# Active monomer RTR-1 derived from the root of Rhodomyrtus tomentosa induces apoptosis in gastric carcinoma cells by inducing ER stress and inhibiting the STAT3 signalling pathway

**DOI:** 10.1101/860130

**Authors:** Xiangqiang Zhang, Jinxia Cheng, Peiyan He, Jinyan Zhu, Zhixian Chen, Shenyu Miao, Guocai Wang, Jianwei Jiang, Yuechun Wang

**Affiliations:** Department of Physiology, Basic Medical College, Jinan University, Guangzhou 510630, China; Department of Biochemistry, Basic Medical College, Jinan University, Guangzhou 510630, China; Department of immunology, Basic Medical College, Jinan University, Guangzhou 510630, China; School of Life Sciences, Guangzhou University, Guangzhou, China; Institute of Traditional Chinese Medicine & Natural Products, College of Pharmacy, Jinan University, Guangzhou 510630, China

## Abstract

**Objective:** *Rhodomyrtus tomentosa*, a flowering plant belonging to the Myrtaceae family, is considered an antitumour substance with versatile biological and pharmacological activities.

And RTR-1 is an active monomer purified from the root of *Rhodomyrtus tomentosa*. However, the detail of the mechanisms of anti-cancer activity of RTR-1 remains to be elucidated and the effect on gastric cancer cells is unknown.

**Methods:** Cell proliferation was determined by MTT and clone formation assay. The effect of RTR-1 on cell cycle and apoptosis was analyzed utilizing flow cytometry, respectively. Moreover, western blotting was used to detect the expression of cell cycle- and apoptosis-related protein.

**Results:** Based on MTT and clone formation assay, we noticed that RTR-1 inhibited the proliferation of gastric carcinoma (BGC823 and SGC7901) cells in a dose- and time-dependent manner. Furthermore, the results of flow cytometry and western blotting showed that RTR-1 induced cell cycle arrest in the G2/M phase through the ATM-Chk2-p53-p21 signalling pathway and induced cell apoptosis by inhibiting the signal transducers and activators of transcription 3 (STAT3) pathway and activating the endoplasmic reticulum stress (ER stress) pathway.

**Conclusion:** Taken together, these results demonstrate that RTR-1 induces cell cycle arrest and promotes apoptosis in gastric carcinoma, indicating its potential application for gastric cancer therapy.

## Introduction

Gastric cancer (GC) remains the second leading cause of cancer deaths worldwide. Each year, approximately 990,000 people are diagnosed with GC worldwide, and approximately 738,000 people die from this disease, [1] making GC the 4^th^ most common cancer by incidence and the 2^nd^ most common cause of cancer death. [2] Although the global incidence of gastric cancer has decreased in recent years, in China, mortality from GC is the highest among all tumours and represents 25% of the mortality caused by gastric cancer worldwide. [3] Although many gratifying achievements are attributable to the use of multidisciplinary approaches for treatment, such as surgery, interventional treatment, biological treatment and so on, drug treatment is still the main treatment for most gastric cancer patients, especially patients with advanced disease.

Despite increased understanding of the biological processes in cancer development, there is still a great need for novel and effective pharmacological strategies for the intervention of many types of cancers. Pharmacological agents that induce apoptosis might be effective against many cancers by inducing death in cancer cells. [4] In fact, a great number of clinically active drugs that are used in cancer therapy are either natural products or are based on natural products. Established plant-derived therapeutics include vinblastine, vincristine, etoposide, teniposide, paclitaxel, doxetaxel, and camptothecin. In addition, numerous plant extracts, including tea polyphenol, resveratrol, ginger extract and soy isoflavones, have been found to demonstrate potential antitumour effects, providing a new direction for the study of new anticancer drugs. [5]

*Rhodomyrtus tomentosa* is a flowering plant belonging to the Myrtaceae family. It is primarily native to Southeast Asian countries, especially the southern parts of Vietnam, China, Japan, Thailand, the Philippines, and Malaysia. It has been used in traditional medicine for the treatment of many diseases, including dysentery and urinary tract infections, and as an antiseptic wash for wounds. In addition, myrtle extracts have been reported to have efficient as antibacterial properties [6] and potent natural antioxidants. [7] Furthermore, a study showed that *Rhodomyrtus tomentosa* has an immunomodulatory effect on innate immune responses [8] and can work as a potential anti-proliferative and apoptosis-inducing agent in HaCaT keratinocytes. [9] In the course of searching for natural anticancer compounds, we evaluated nine compounds for their anti-proliferative action in BGC823, SGC7901, SK-Mel-110 and SMMC7721 cells. RTR-1 was found to induce potent cytotoxic effects in four strains of tumour cells. Eight compounds induced potent cytotoxic effects in some cells. By consulting the literature, we found that RTR-1 is a newly isolated compound from *Rhodomyrtus tomentosa*. Therefore, we chose RTR-1 as the research subject to explore the molecular mechanism by which it inhibits the growth of gastric cancer cells.

In this study, we found that RTR-1 blocks cell cycle progression at the G2/M phase in a ROS-dependent manner. Moreover, RTR-1 also induces caspase-regulated apoptotic cell death by activating ER stress and inhibiting the STAT3 signalling pathway. Our study suggests that RTR-1 may be a new source of anticancer compounds.

## Material and methods

### General experimental procedures

Optical rotations were recorded on a JASCO P-1030 automatic digital polarimeter, and UV spectra were recorded with a JASCO V-550 UV/VIS spectrophotometer. IR spectra were determined using a JASCO FT/IR-480 plus FT-IR spectrometer. HRESIMS data were determined by an Agilent 6210 ESI/TOF mass spectrometer. NMR spectra were obtained by a Bruker AV-400 spectrometer with TMS as an internal standard. Preparative HPLC was performed using a Varian chromatograph equipped with a C18 reversed-phase column (Cosmosil, 5 µm, 10 mm × 250 mm). Analytical HPLC was performed using a Waters chromatograph equipped with a C18 reversed phase column (Cosmosil, 5 µm, 4.6 mm × 250 mm). Silica gel (200–300 mesh; Qingdao Marine Chemical, Inc.), ODS silica gel (50 µm; YMC), and Sephadex LH-20 (Pharmacia) were used for column chromatography experiments. Silica gel GF254 plates (Yantai Chemical Industry Research Institute, Yantai, China) were used for thin-layer chromatography (TLC).

### Materials

The dried roots of *R. tomentosa* were purchased in Guangzhou, Guangdong Province, China, in March 2013. The plant was authenticated by Zhenqiu Mai, the senior engineer of a medicinal materials company in Guangdong Province. A voucher specimen (20130330) was deposited in the Institute of Traditional Chinese Medicine and Natural Products of Jinan University.

### Extraction and isolation

The dried roots of *R. tomentosa* (25.0 kg) were pulverized and extracted with 95% aqueous ethanol (100 L) at 50 °C three times. The ethanol extract was concentrated in vacuo to obtain a crude extract (1.6 kg). The crude extract was suspended in water and partitioned with petroleum ether (2.5 g) and ethyl acetate (651.3 g). The ethyl acetate extract was subjected to silica gel column chromatography using a cyclohexane/ethyl acetate system (100:0 to 0:100, v:v) in eight fractions (Fr. A-H). Moreover, Fr. D was eluted by chromatography with a chloroform/methanol gradient on a silica gel column, which yielded compound RTR-9 (240.5 mg) and RTR-10 (3.3 mg). Additionally, Fr. G was further separated by silica gel column chromatography with chloroform/methanol (100:0 to 0:100, v:v) and was purified by a Sephadex LH-20 (CHCl_3_/MeOH, 50:50, v/v) column and preparative HPLC with MeOH-H_2_O, which yielded compound RTR-1 (124.5 mg), RTR-2 (30.6 mg), RTR-3 (42.6 mg), RTR-4 (26.7 g), RTR-5 (72.8 mg), RTR-6 (81.4 mg), RTR-7 (8.4 mg), RTR-8 (24.1 mg), RTR-11 (3.4 mg), RTR-14 (0.9 g), RTR-15 (6.1 mg), RTR-16 (2.1 mg), RTR-17 (1.4 mg), RTR-18 (542.5 mg), and RTR-19 (9.0 mg). Moreover, the chemical structure of RTR-1, RTR-2, RTR-3, RTR-4, RTR5, RTR-6, RTR-8, RTR-9, RTR-17 are showed in Figure 1A.

**Fig 1.**
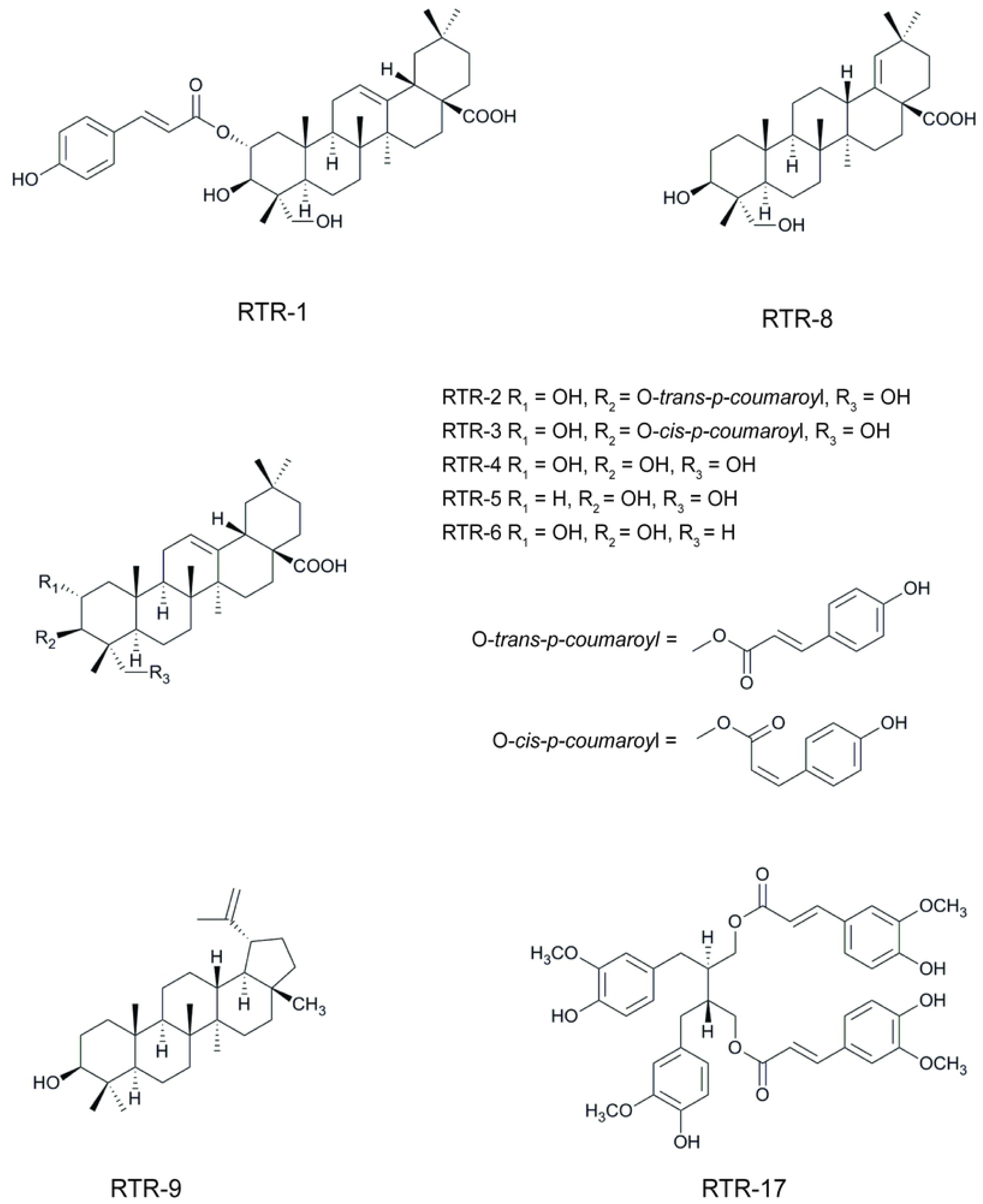
The chemical structure of RTR-1, RTR-2, RTR-3, RTR-4, RTR-5, RTR-6, RTR-8, RTR-9, and RTR-17.

The following data were obtained on RTR-1: white powder; [α]25 D +10.7 (c 0.64, CH_3_OH), HRESIMS m/z 657.3762 [M+Na]+ (calcd for C_39_H_54_O_7_Na, 657.3762); UV (CH_3_OH) λmax 227, 313 nm; IR (KBr) λmax 3312, 2946, 1698, 1631, 1604, 1515, 1455, 1270, 1170, 1048, and 831 cm-1; 1H NMR (400 MHz, Pyr-d5) δH: 7.93 (1H, d, J = 15.6 Hz, H-3’), 7.51 (2H, d, J = 8.4 Hz, H-5’,9’), 7.13 (2H, d, J = 8.4 Hz, H-6’, 8’), 6.53 (1H, d, J = 15.6 Hz, H-2’), 5.79 (1H, ddd, J = 12.4, 10.0, 4.4 Hz, H-2), 5.50 (1H, br s, H-12), 4.49 (1H, d, J = 10.0 Hz, H-3), 1.26 (3H, s, H-27), 1.17 (3H, s, H-25), 1.09 (3H, s, H-24), 1.03 (3H, s, H-26), 0.98 (3H, s, H-30), and 0.91 (3H, s, H-29); 13C NMR (100 MHz, Pyr-d5) δC: 44.9(C-1), 74.0 (C-2), 74.3 (C-3), 44.6 (C-4), 47.8 (C-5), 18.8 (C-6), 33.1 (C-7), 40.1 (C-8), 48.4 (C-9), 38.9 (C-10), 24.0 (C-11), 122.7 (C-12), 145.3 (C-13), 42.6 (C-14), 28.6 (C-15), 24.3 (C-16), 46.9 (C-17), 42.3 (C-18), 46.7 (C-19), 31.3 (C-20), 34.5 (C-21), 33.5 (C-22), 65.9 (C-23), 14.7 (C-24), 17.5 (C-25), 17.8 (C-26), 26.5 (C-27), 180.5 (C-28), 33.6 (C-29), 24.1 (C-30), 167.9 (C-1’), 116.4 (C-2’), 144.9 (C-3’), 126.5 (C-4’), 130.8 (C-5’), 117.1 (C-6’), 161.6 (C-7’), 117.1 (C-8’), and 130.8 (C-9’).

### Cell lines and reagents

For this study, BGC823 and SGC7901 cells (human gastric cancer cell lines) and LO2 cells (a human hepatic cell line) were obtained from the American Type Culture Collection (Rockville, MD) and were cultivated in DMEM supplemented with 10% foetal bovine serum in a 5% CO2 humidified atmosphere at 37°C. Foetal bovine serum, DMEM, trypsin and EDTA were purchased from Gibco; MTT and DMSO were purchased from Sigma; annexin V-fluorescein isothiocyanate (FITC) and propidium iodide (PI) were purchased from BD Biosciences. Antibodies against P21, P53, c-IAP1, c-IAP2, GAPDH, Bcl-2, and Bcl-xL were purchased from Santa Cruz Biotechnology (Santa Cruz, CA). Mcl-1, Bax, Caspase-3, cleaved Caspase-3, Caspase-9, PARP, stat3, p-Stat3, ATM, p-ATM, chk2, p-chk2 were purchased from Cell Signaling Technology (Danvers, MA). RTR-1, derived from *Rhodomyrtus tomentosa* root and provided by the Institute of Traditional Chinese Medicine and Natural Products of Jinan University, was dissolved in dimethyl sulfoxide (DMSO; Sigma, USA) at 100 mmol/L.

### MTT Assay

Cell growth and viability were measured using an MTT assay. The cells in the logarithmic growth phase were seeded in 96-well plates at a density of 5.0×10^3^ cells per well and incubated overnight. Then, the cells were treated with different concentrations of RTR-1 (0.78, 1.56, 3.12, 6.25, 12.5, 25 and 50 μmol/L), and DMSO (0.04%) was used as the vehicle control (5 repeated wells were set up in each group). The cells were incubated between 24 h and 72 h in standard culture conditions. Then, 10 μL of MTT (5 mg/mL) was added to each well and the cells were incubated for an additional 4-6 h. The supernatants were then carefully discarded, and 100 μL of DMSO was added; the cells were then shaken vigorously for 5 min to dissolve the purple precipitate. A microplate reader (Bio-Rad Laboratories, Hercules, CA, USA) was used to measure the absorbance at a wavelength of 570 nm. The following formula was used to calculate the inhibition rate of cell proliferation: cell inhibitory rate = (1-A_treatment_/A_blank_) × 100%. Each measurement was performed in triplicate, and each experiment was conducted at least three times.

### Colony formation assays

Cells in the logarithmic growth phase were seeded at a density of 500 cells per well in 6-well tissue culture plates and incubated overnight. Then, the cells were treated with different concentrations of silibinin (2, 4, 8 μmol/L), and DMSO (0.04%) was used as the vehicle control. The cells were incubated for 14 days under standard culture conditions. At the end of the incubation period, the cells were fixed with methanol:acetic acid = 3:1 and stained with 0.1% (w/v) crystal violet. Megascopic cell colonies were photographed with a Canon camera. Each measurement was taken in triplicate, and each experiment was conducted at least three times.

### Cellular ROS production

*C*ells were seeded in six-well plates with 2 mL in each well. The cells were treated with either RTR-1 for the indicated times, with various concentrations of RTR-1 for 24 h or pre-incubated with 5 mmol/L NAC followed by treatment with 40 μmol/L RTR-1 for 24 h and then incubated with 5 μmol/L (final concentration) CM-H2DCF-DA in the dark for 30 min at 37°C. After being washed twice with PBS at 4°C, the cells were centrifuged and resuspended in PBS. The levels of intracellular ROS were detected by flow cytometry with a FACSCalibur system and CellQuest PRO analysis software.

### Cell cycle distribution

Cells in the logarithmic growth phase were seeded at a density of 1.0×10^5^ cells per well in 6-well tissue culture plates. Then, the cells were treated with different concentrations of RTR-1 (10, 20, or 40 μmol/L), and DMSO (0.04%) was used as the vehicle control. The cells were washed twice with cold PBS and harvested by trypsinization. The cells were washed twice with cold PBS and then fixed in 70% ethanol overnight at 4°C. Then, the cells were washed three times with PBS and incubated with PI (500 g/L) and RNase for 0.5 h in the dark. The samples were immediately analysed using a flow cytometer. A total of 10,000 cells was analysed in each sample.

### Annexin V-FITC/PI double-staining assay

Cells in the logarithmic growth phase were seeded at a density of 2.0×10^5^ cells per well in 6-well plates. Then, the cells were treated with various concentrations of RTR-1 (10, 20, or 40 μmol/L), and DMSO (0.04%) was used as the vehicle control. After 24 h of incubation, both the floated and attached cells were collected. The cells were washed twice with ice-cold PBS, resuspended in 200 μL of binding buffer, and incubated with 10 μL of annexin V-FITC and 5 μL of PI for 15 min at room temperature in the dark. Then, a flow cytometer was used to analyse the percentage of cells in early and late apoptosis. A total of 10,000 cells was analysed in each sample.

### Hoechst 33342 staining

Cells in the logarithmic growth phase were seeded at a density of 2.0×10^5^ cells per well in 6-well plates. Then, the cells were treated with various concentrations of RTR-1 (10, 20, or 40 µmol/L), and DMSO (0.04%) was used as the vehicle control. After 24 h of incubation, the cells were stained with 2 μg/mL Hoechst 33342 dye for 5 min. The cells were then washed twice with PBS. The cells were detected at an excitation wavelength of 350 nm and emission wavelength of 460 nm by fluorescence microscopy.

### Western blot analysis

Cells were seeded at a density of 2.0×10^5^ per well in 6-well plates. Then, the cells were treated with various concentrations of RTR-1 (10, 20, and 40 µmol/L), and DMSO (0.04%) was used as the vehicle control. After 24 h of incubation, the cells were collected and washed twice with ice-cold PBS. Then, the cells were lysed in buffer on ice for 15 min and centrifuged at 12,000 × g for 10 min at 4 °C. The cell proteins were extracted, and the protein concentration was determined using a BCA assay. A total of 25 µg of protein was denatured by the addition of 6× reducing sample buffer after incubation for 10 min at 100 °C, and then, the protein samples (25 μg) were loaded in each well of an 8% −15% polyacrylamide gel, separated by SDS-PAGE and transferred to a polyvinylidene fluoride (PVDF) membrane (Millipore, Bedford, MA, USA). After the membranes were blocked with 5% skimmed milk, the membranes were incubated overnight at 4 °C with the appropriate antibody. Then, the membranes were washed three times for 10 min each time with TBST (10 mM Tris, 100 mM NaCl, and 0.1% Tween 20) and incubated for 2 h at room temperature with a secondary antibody conjugated to horseradish peroxidase. After the membrane was washed three times in TBST, the bound antibody complex was detected using an ECL chemiluminescence reagent and XAR film (Kodak) as described previously.

### Statistical analyses

The results are expressed as the mean ± standard error (SE) of 3 independent experiments. Student’s t-tests were used to determine the significance between the control and test groups. P<0.05 was considered significant.

## Results

### Biological activity

To verify the potential anticancer properties of the RTR compounds isolated from *Rhodomyrtus tomentosa*, we first tested whether the compounds could inhibit the proliferation of various cancer cells. To this end, gastric cancer (BGC823 and SGC7901) cells, hepatocellular cancer (SMMC7721) cells and malignant melanoma (SK-Mel-110) cells were treated with increasing concentrations of the RTR compounds, and the extent of the cell proliferation was determined with an MTT assay. As shown in Figure 2A-D, most RTR compounds could partially inhibit cell proliferation in a dose-dependent manner in cancer cell lines. Furthermore, the inhibition concentration (IC50) values after 48 h RTR compound exposure on BGC823, SGC7901, SMMC7721 and SK-Mel-110 cells are shown in Figure 2E. More interestingly, the IC50 values of RTR-1 in the BGC823, SGC7901, SMMC7721 and SK-Mel-110 cells were 15.43 ± 0.47, 16.80 ± 0.4, 20.89 ± 0.84 and 12.66 ± 0.3 μmol/L, respectively. These data suggest that RTR-1 may be a new source of anticancer compounds.

**Fig 2.**
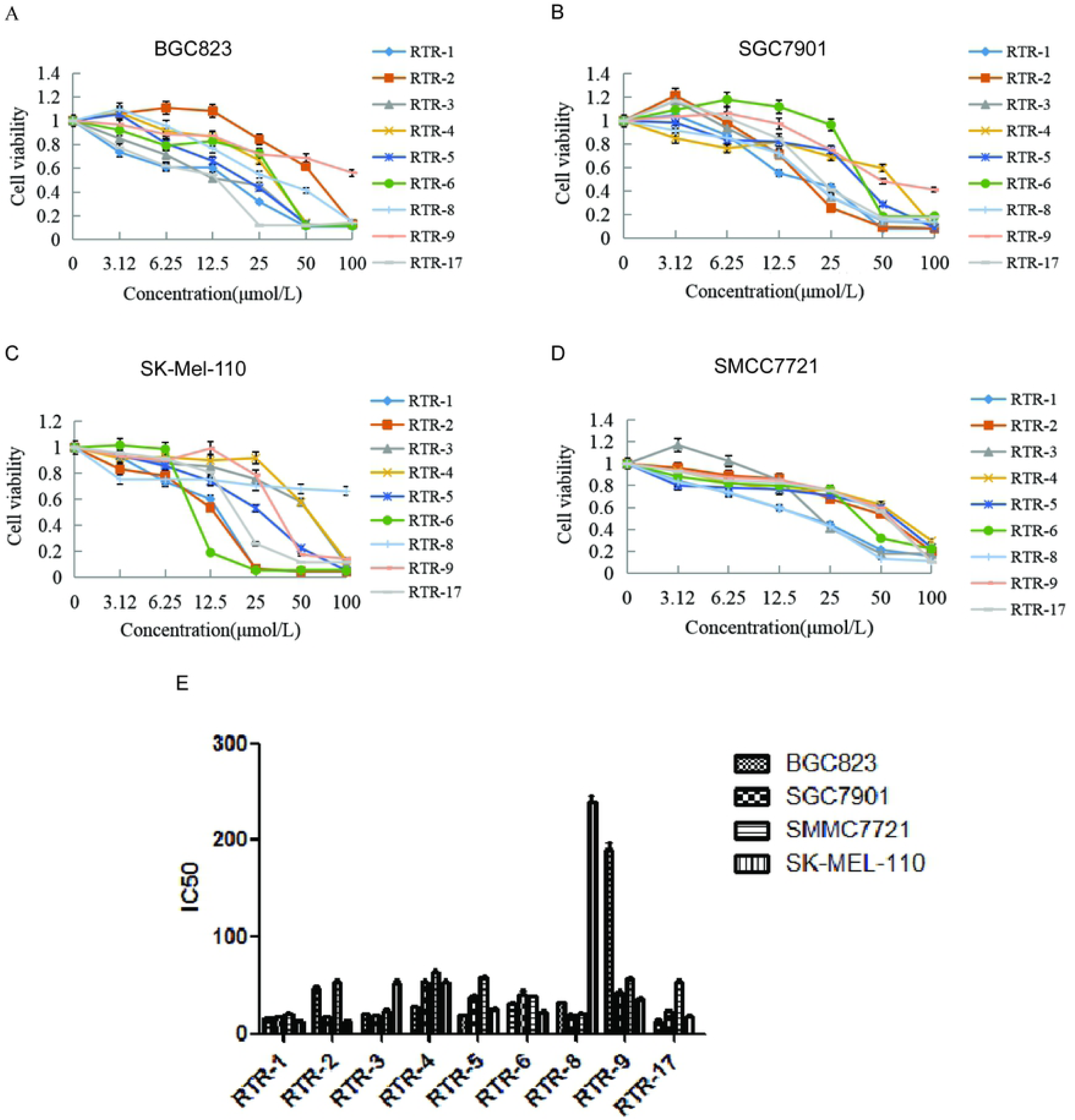
Inhibitory activities of RTRs in some cancer cell lines. (A) Proliferative effects of the RTR compounds on the BGC823 cells at 48 h. (B) Proliferative effects of the RTRs on the SGC7901 cells at 48 h. (C) Proliferative effects of RTRs on the SK-Mel-110 at 48 h. (D) Proliferative effects of the RTRs on the SMMC7721 cells at 48 h. Each value represents three independent experiments. (E) The IC50 values of RTR compounds for the BGC823, SGC7901, SK-Mel-110 and SMMC7721 cells at 48 h.

### Growth inhibition effects of RTR-1 on gastric cancer cells and LO2 cells

To investigate the anticancer effect of RTR-1 on gastric cancer proliferation and viability, we selected normal liver cancer (LO2) cells and cells from two human gastric cancer cell lines (BGC823 and SGC7901) and treated them with various concentrations of RTR-1 for 24 h, 48 h and 72 h (Figure 3A-C). Our results showed that RTR-1 treatment decreased the viability of the BGC823 and SGC7901 cells in a dose- and time-dependent manner. Interestingly, the effect of RTR-1 on LO2 cells was not as notable. In addition, the IC50 values of RTR-1 for the LO2 cells at 24 h, 48 h and 72 h were 154.36±5.26, 70±3.33, and 46.82 ± 4.45 μmol/L, respectively. The IC50 values of RTR-1 on the BGC823 cells at 24 h, 48 h and 72 h were 32.47±1.23, 15.43±0.47, and 6.40±0.03 mol/L, respectively. The IC50 values of RTR-1 on the SGC7901 cells at 24 h, 48 h and 72 h were 32.10±1.09, 16.80±0.4, 8.38±0.56 μmol/L, respectively (Figure 3D). Furthermore, the colony formation assays showed that RTR-1 could dramatically inhibit colony formation in a concentration-dependent manner, as indicated by fewer and small colonies in the drug-treated group (Figure 3E). Taken together, these results demonstrate that RTR-1 exhibits significant anticancer activity by inhibiting cell proliferation and viability.

**Fig 3.**
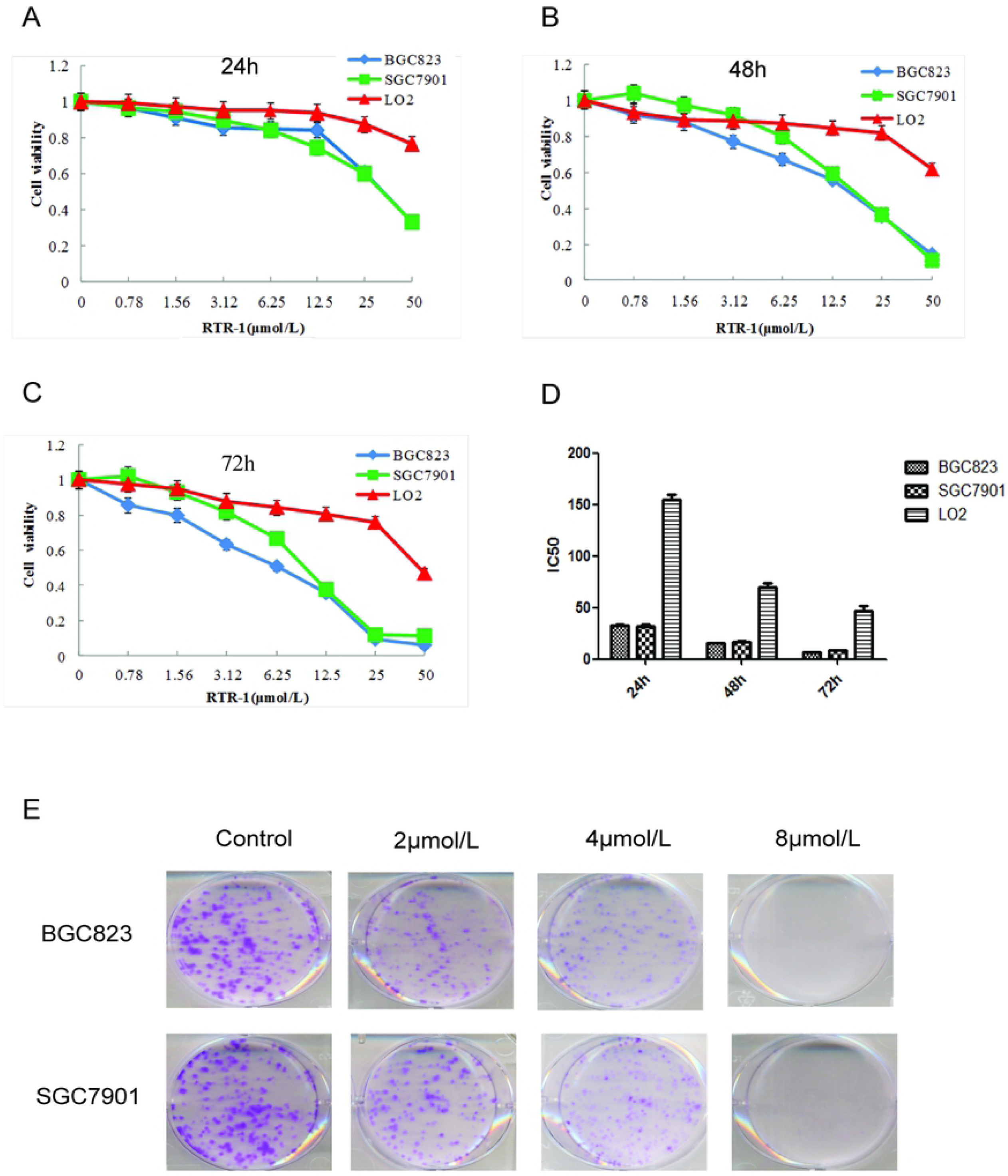
Inhibitory activities of RTR-1 in the BGC823, SGC7901 and LO2 cells. (A) IC50 values of RTR-1 for the BGC823, SGC7901 and LO2 cells in vitro. (B, C, D) Proliferative effects of RTR-1 on the BGC823, SGC7901, and LO2 cells. Inhibition of cell proliferation was determined with MTT assays, and the IC50 value was calculated as described in the materials and methods. (E) BGC823 and SGC7901 cells were treated with different concentrations of RTR-1 for 14 days. Each value represents three independent experiments.

### TR-1 induces cell cycle arrest through the ATM-Chk2-p53-p21 signalling pathway

Cell proliferation inhibition is often strongly associated with changes in cell cycle progression; [10] therefore, we evaluated the distribution of BGC823 and SGC7901 cells in the cell cycle. After treatment with RTR-1 for 24 h, the cell cycle distribution changed significantly, with a remarkable increase in the cell population in the G2/M phase and a decrease in the cell population in the S phase (Figure 4A). For example, the fraction of the BGC823 cells in the G2/M phase increased from 13.5% (DMSO treated) to 20.8%, 24.5% and 32.1% for cells treated with 10, 20, and 40 μM RTR-1, respectively. In summary, the cell cycle distribution results indicate that the cell cycle is mainly arrested in the G2/M phase.

**Fig 4.**
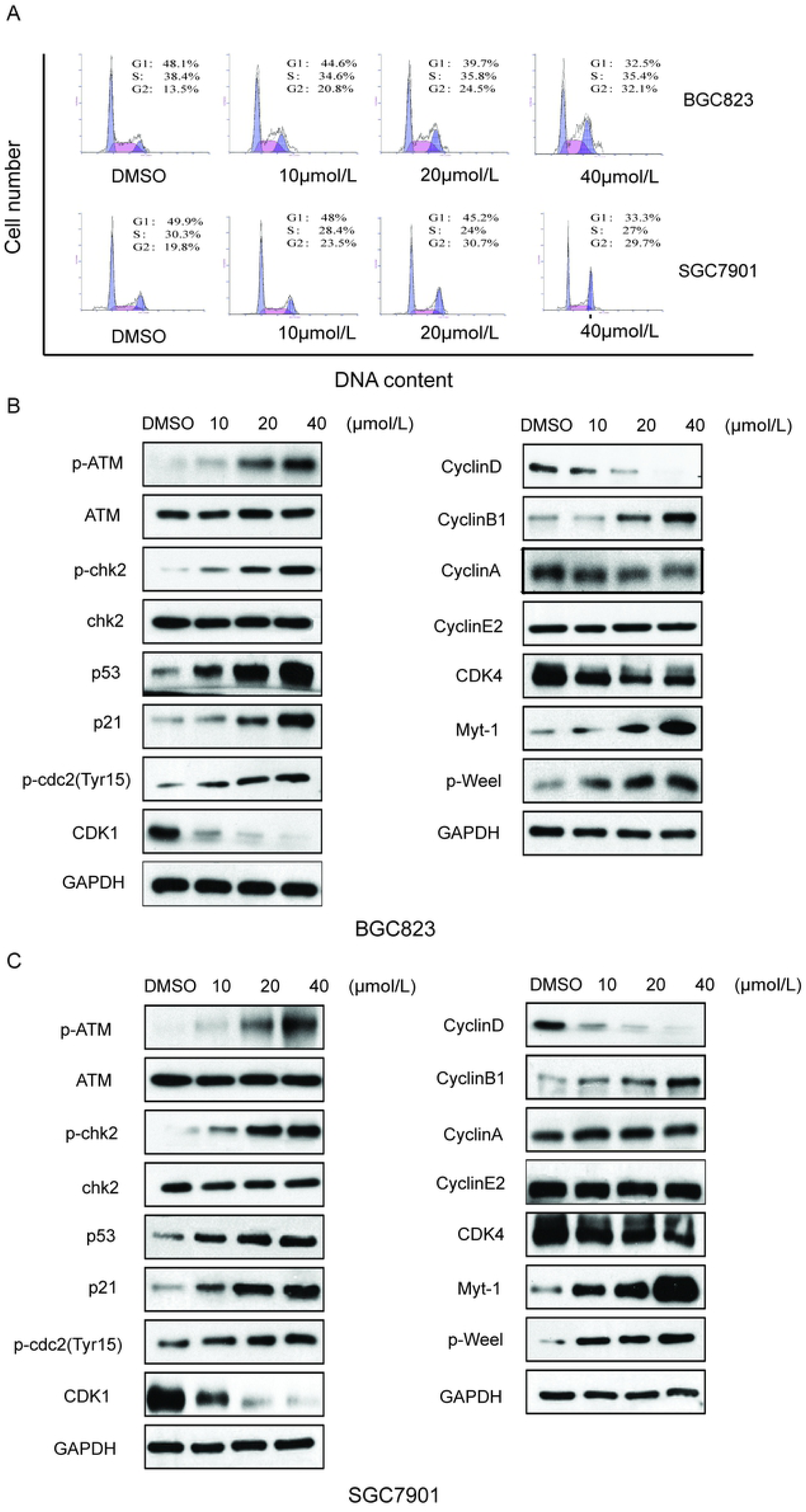
Effect of RTR-1 on the induction of cell cycle arrest in the BGC823 and SGC7901 cells. (A) BGC823 and SGC7901 cells were treated with various concentrations of RTR-1 for 24 h. The expression of the cell cycle-associated proteins in the BGC823 cells (B) and the SGC7901 cells (C) was analysed by Western blotting. Data represent one of three experiments that yielded similar results.

Cyclins and CDK complexes are of great significance in regulating cell cycle progression, with different CDKs and cyclins governing different phases of the cell cycle. In gastric cells, the cyclin B/cdc2 complex has an important function in controlling the G2/M transition. However, the function of the cyclin B/cdc2 complex can be reversed by p21 and Myt-1, as Myt-1 phosphorylates Tyr15 and Thr14 of CDK1 (cdc2) and inhibits its activity. We used a Western blot analysis to determine the expression of cell cycle-related proteins, namely, cyclins, CDKs, CKIs, Myt-1 and p-Weel. As shown in Figure 4B and C, the expression levels of Myt-1, cyclin B1, P-cdc2 (Tyr15) and p-Weel were markedly increased, while cyclin D1, CDK1 and CDK4 were noticeably reduced, and the expression levels of cyclin E and cyclin A remained unchanged in the RTR-1-treated groups compared to the levels in the control groups. It has been reported that, in response to DNA damage, cells activate a signalling network to arrest the cell cycle and facilitate DNA repair, which inhibits cell proliferation. [11] To determine how RTR-1 induced cell cycle arrest in the G2/M phase, we examined the expression levels of the proteins involved in the ATM-Chk2-p53-p21 signalling pathways using Western blotting. We found that the expression levels of p-ATM, p-chk2, p53, and p21 were increased (Figure 4B and C). Thus, these data further demonstrate that RTR-1 induces cell cycle arrest in the G2/M phase through the ATM-Chk2-p53-p21 signalling pathways in BGC823 and SGC7901 cells.

Overall, these results indicate that RTR-1 inhibits cell proliferation by inducing cell cycle arrest in the G2/M phase through the ATM-Chk2-p53-p21 signalling pathway in BGC823 and SGC7901 cells.

### RTR-1 promotes apoptosis via the caspase pathway

Phosphatidylserine (PS) translocation to the cell surface is an indicator of early apoptosis. To confirm whether the growth inhibition of RTR-1 was caused by apoptosis in vitro, we first used an annexin V-FITC/PI double-staining assay to detect the number of apoptotic BGC823 and SGC7901 cells that had been treated with 0.08% DMSO or 10, 20 and 40 µmol/L RTR-1 for 24 h. As illustrated in Figure 5A, the percentage of apoptotic cells increased with increasing RTR-1 concentrations. Moreover, the apoptosis rate reached 41.8% in the RTR-1 treated BGC823 cell, compared to 1.2% for the control cells, while it reached 46.2% for the RTR-1 treated SGC7901 cells compared to 2.2% for the control cells. Furthermore, Hoechst 33342 staining showed that the BGC823 and SGC7901 cells treated with increasing RTR-1 concentrations showed significant apoptotic changes; for example, the cytoskeletal structures were damaged, and the cells became round and shrunken with nuclear fragmentation and aggregation (Figure 5B). In addition, we were further inspired to explore how RTR-1 is involved in the activation of apoptosis. Caspase-9 is a major caspase member that mediates intrinsic apoptosis, while caspase-3 is one of the key effectors in the execution of the downstream apoptotic pathway. [12] After treatment with RTR-1 for 24 h, upregulated levels of cleaved caspase-3 and downregulated levels of PARP, caspase-3 and caspase-9 were observed in the BGC823 and SGC7901 cells (Figure 5C). Taken together, these results indicate that RTR-1 inhibits cell proliferation by inducing apoptosis through the caspase pathway in BGC823 and SGC7901 cells.

**Fig 5.**
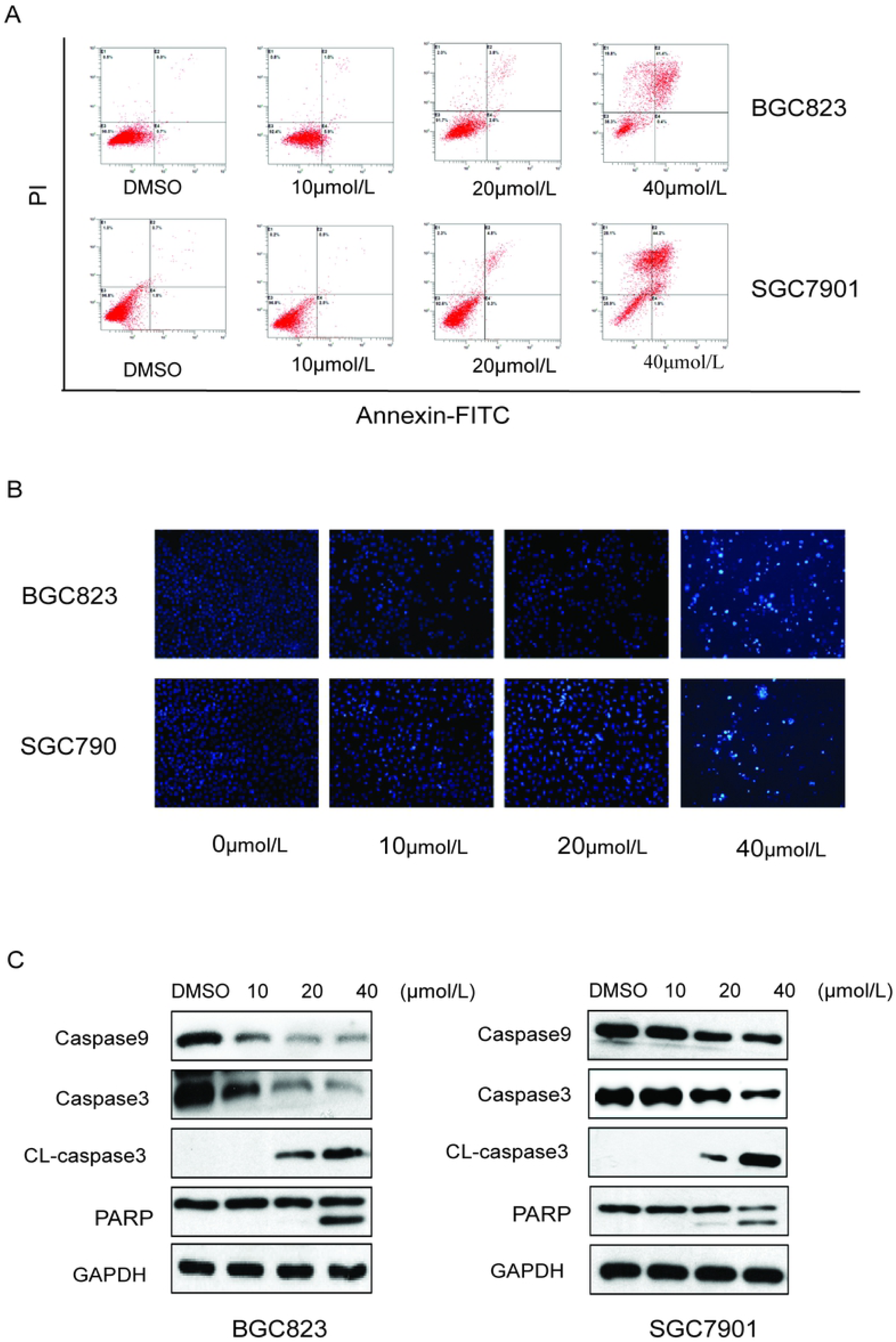
Effect of RTR-1 on the induction of apoptosis in the BGC823 and SGC7901 cells. (A) BGC823 and SGC7901 cells were treated with various concentrations of RTR-1 for 24 h and stained with annexin V-FITC and PI to distinguish and quantitatively determine the percentages of apoptotic cells. (B) BGC823 and SGC7901 cells were treated with various concentrations of RTR-1 for 24 h and analysed with Hoechst 33342 staining. (C) The expression of apoptosis-related proteins in the BGC823 and SGC7901 cells was analysed by Western blotting. Data represent one of three experiments that yielded similar results.

### Effects of RTR-1 on the expression of IAPs and Bcl-2 family proteins

The generation of reactive oxygen species (ROS) has been reported to play an important role in the cell cycle. [13] In this study, we found that the level of reactive oxygen species increased when BGC823 and SGC7901 cells were treated with RTR-1 in a dose- and time-dependent manner (Figure 6A). Additionally, the Bcl-2 family of proteins has great significance with regard to apoptosis. Bcl-2 family proteins, including the pro-apoptotic proteins Bax and Bad and the anti-apoptotic proteins Bcl-2, Bcl-xl and Mcl-1, were also investigated. As shown in Figure 6B, RTR-1 treatment significantly reduced the expression of Mcl-1 and Bcl-2 and enhanced the expression of Bax and Bad but did not change the expression of Bcl-xl. These results suggest that the apoptosis induced by RTR-1 occurs through the mitochondria-mediated pathway in BGC823 and SGC7901 cells. Furthermore, the apoptosis induced by overexpression of pro-caspases 3, 7 or 9 can also be suppressed by the co-expression of XIAP, c-IAP1, c-IAP2 and Survivin. In this study, we found that the expression of XIAP, c-IAP1 and c-IAP2 decreased and that the expression of survivin decreased in the BGC823 cells, while the expression of survivin remained unchanged in the SGC7901 cells (Figure 6C).

**Fig 6.**
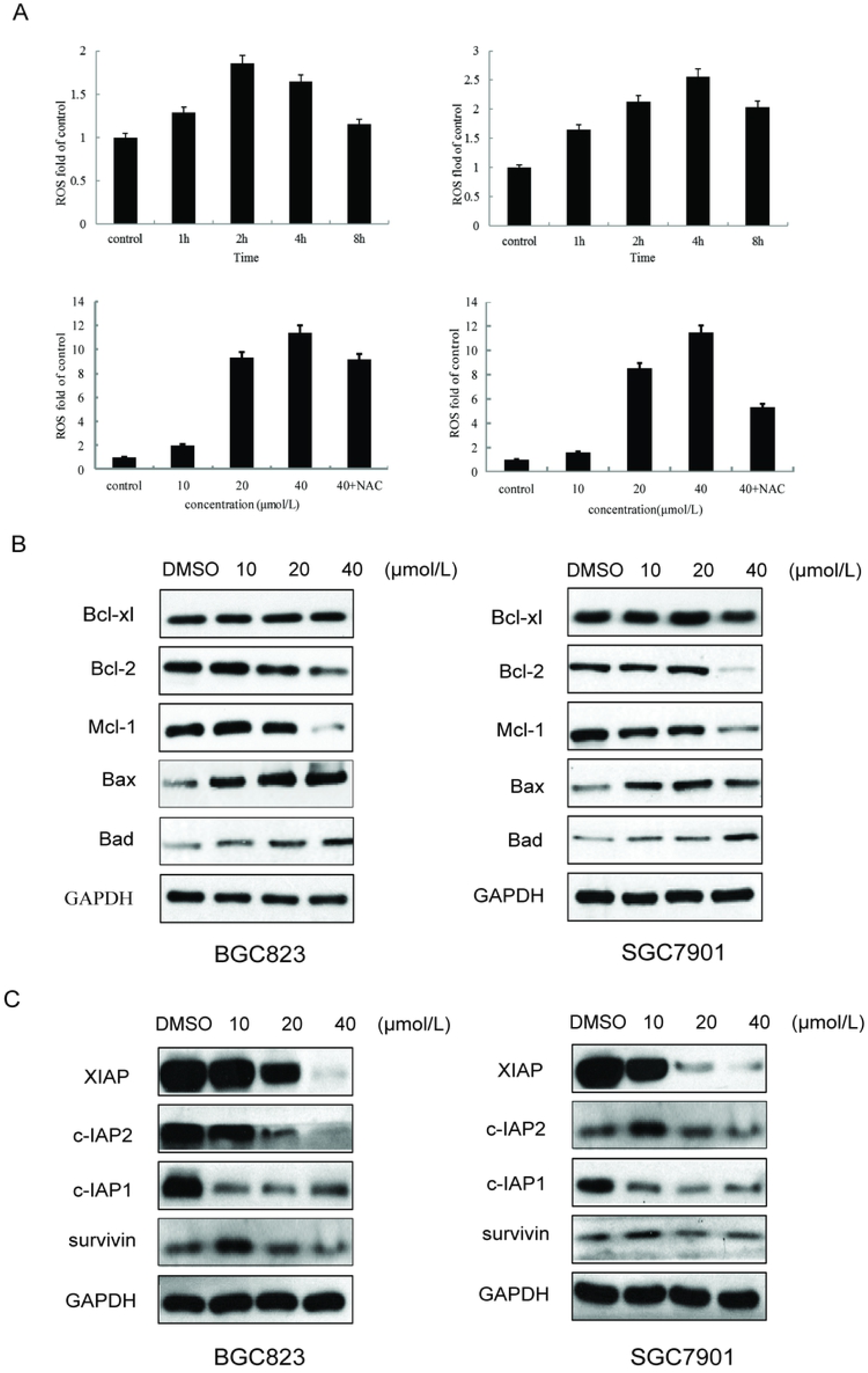
Effect of RTR-1 on the expression of IAP and Bcl-2 family proteins. (A) BGC823 and SGC7901 cells were treated with 20 μmol/L RTR-1 for different time periods (hours) and treated with different concentrations of RTR-1 for 24 h in the presence or absence of NAC. (B) Bcl-2 family proteins were analysed by Western blotting. (C) IAP family proteins were analysed by Western blotting. Data represent one of three experiments that yielded similar results.

These findings suggest that ROS generation is the key regulator of RTR-1-induced cell cycle arrest and apoptosis induced by RTR-1 through the mitochondria-mediated and IAP pathways in BGC823 and SGC7901 cells.

### RTR-1 induces ER stress and inhibits the STAT3 pathway

It has been found that endoplasmic reticulum (ER) stress can trigger apoptotic signalling. To investigate whether RTR-1-induced apoptosis is mediated by ER stress, we used Western blot analysis to determine the levels of ER stress-associated proteins in the BGC823 and SGC7901 cells. After treatment with different doses of RTR-1 for 24 h, the levels of IRE1α, Erol-Lα, CHOP, PERK and BiP were upregulated, while there was less effect on the expression of PDI (Figure 7A). Thus, these data demonstrate that RTR-1 promotes apoptosis by inducing ER stress in BGC823 and SGC7901 cells.

**Fig 7.**
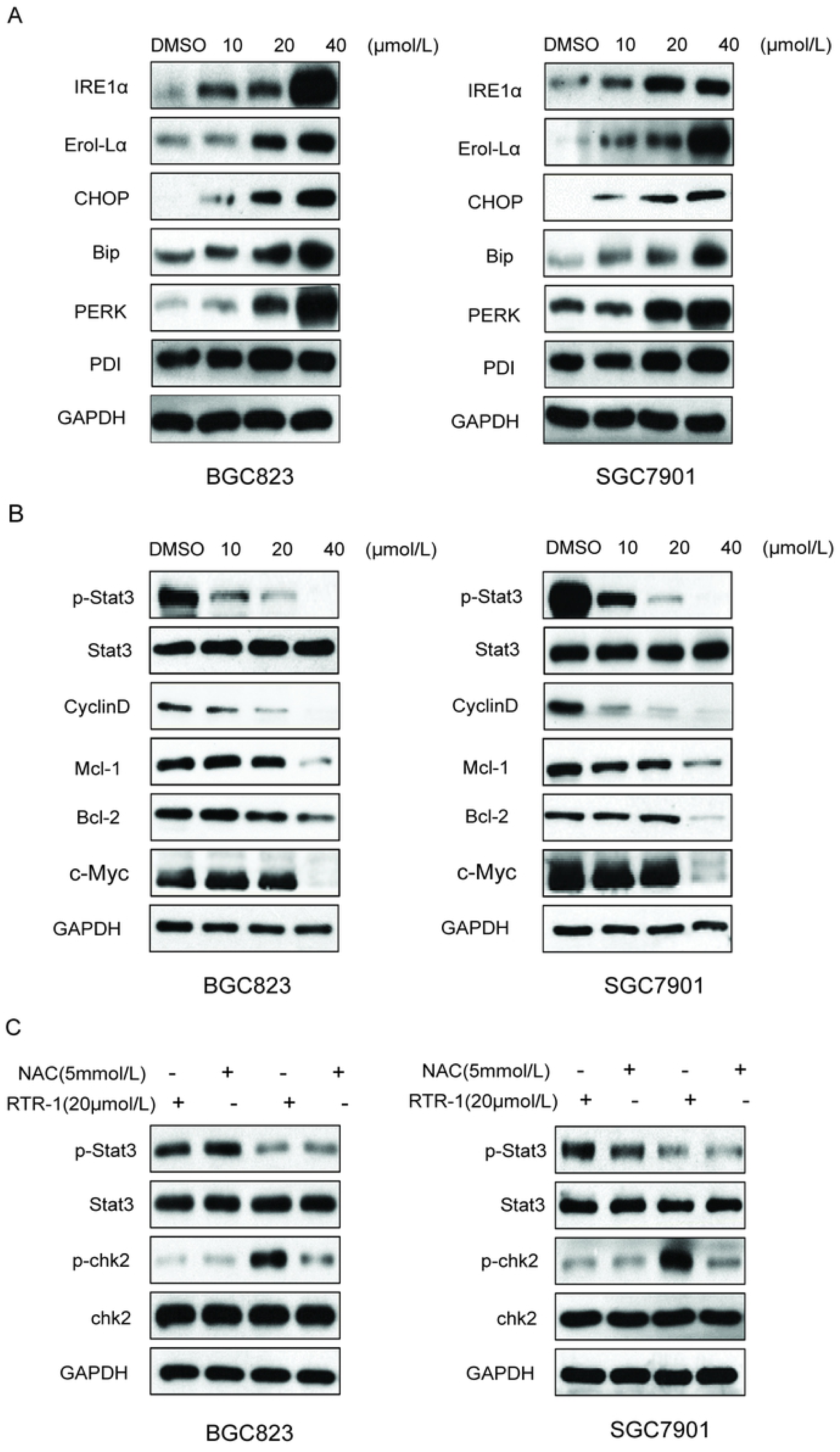
Effect of RTR-1 on the ER stress and STAT3 pathways in the BGC823 and SGC7901 cells was analysed by Western blotting. (A) ERS-related proteins were analysed by Western blotting. (B) STAT3-related proteins were analysed by Western blotting. (C) The effect of ROS on the Stat3 signal pathway and cell cycle-associated proteins in the BGC823 and SGC7901 cells was analysed by Western blotting.

In addition, signal transducer and activator of transcription 3 (STAT3), an oncogenic transcription factor, promotes tumorigenesis by regulating the expression of various target genes, including cell cycle regulators, angiogenic factors and anti-apoptosis genes. [14] To study whether RTR-1 modulated STAT3 activation, the BGC823 and SGC7901 cells were treated with different concentrations of RTR-1, and then, STAT3 and STAT3 phosphorylation were examined by Western blotting. We found that p-STAT3 in the BGC823 and SGC7901 cells was substantially reduced upon RTR-1 treatment (Figure 7B). Moreover, we analysed the effect of RTR-1 on some proteins that mediate anti-apoptotic signals downstream of STAT3 activation, namely, cyclin D1, c-Myc, Bcl-2 and Mcl-1. As shown in Fig. 8B, RTR-1 strongly inhibited cyclin D1, c-Myc, Bcl-2 and Mcl-1 protein expression in the BGC823 and SGC7901 cells. Therefore, RTR-1 effectively inhibits the STAT3 pathway in BGC823 and SGC7901 cells.

**Fig 8.**
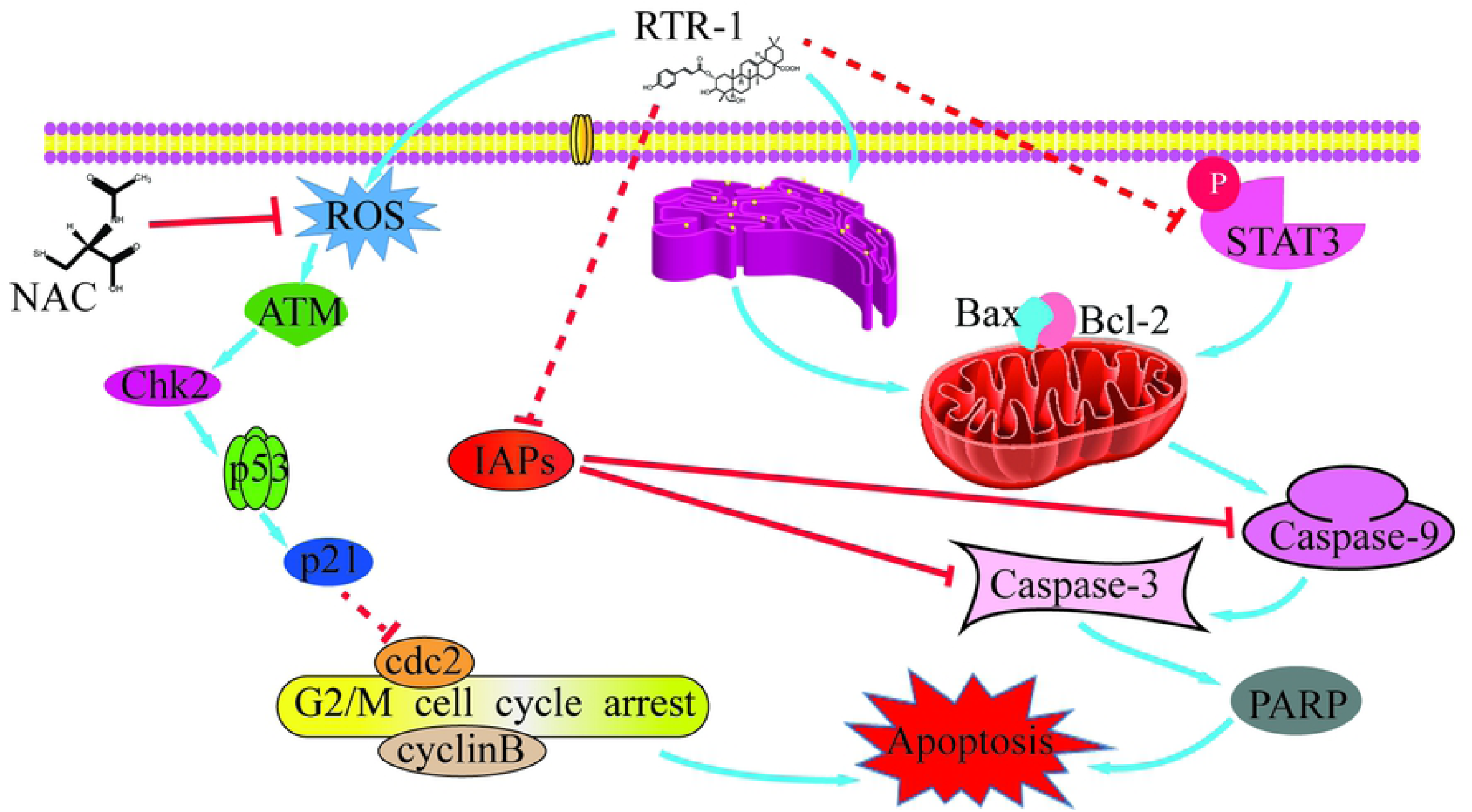
Schematic model for the ability of RTR-1 to induce cell cycle arrest and apoptosis.

Furthermore, ROS may play roles in cytokine activation of JAK kinases as well as STAT3 and STAT5 expression and the cell cycle. [15] To further confirm the relationship between ROS and STAT3 and the cell cycle, the expression of p-STAT3 and p-chk2 was examined by Western blot analysis after N-acetyl-L-cysteine (NAC) pretreatment. Our results show that pretreatment with NAC significantly reversed the RTR-1-induced increase in p-chk2 in the BGC823 and SGC7901 cells. However, the expression of p-STAT3 was less affected after pretreatment with NAC in the BGC823 and SGC7901 cells (Figure 7C). These data further demonstrate that ROS play an essential role in inducing cell cycle arrest in BGC823 and SGC7901 cells.

Thus, our results indicate that RTR-1 inhibits gastric cancer cell growth via the promotion of apoptosis by inducing ER stress and inhibiting the STAT3 signalling pathway.

## Discussion

Gastric cancer is associated with high morbidity and mortality worldwide. Chemotherapy is a cornerstone of the treatments for gastric cancer; however, less than 60% of patients receive salvage therapy in clinical practice. Moreover, gastric cancer has poor outcomes with regard to both surgical resection and chemotherapy. [16] Therefore, the search for an effective chemotherapeutic drug that induces minimal toxicity has become an important part of the search for gastric cancer treatments. Additionally, studies on the activities of compound extracts from myrtle leaves, stems and flowers have become the focus of research. It has been discovered that compound extracts from myrtle leaves, stems and flowers exhibit antioxidant effects [17] and potential antibiotic and anti-infective effects. ^[18]^ However, it remains unclear whether RTR-1, a compound recently isolated from *Rhodomyrtus tomentosa*, can inhibit the proliferation of gastric cancer cells.

In this study, we first investigated the inhibitory effect of RTR-1 on the proliferation of BGC823, SGC7901 and LO2 cells by MTT assay and found that RTR-1 had a relatively weak inhibitory effect on the proliferation of the LO2 cells compared to the significant time- and dose-dependent growth inhibition effects it had on the BGC823 and SGC7901 cells. The inhibition concentration (IC50) values of RTR-1 on the LO2 cells at 24 h, 48 h and 72 h were 154.36±5.26, 70±3.33, and 46.82 ± 4.45 μmol/L, respectively. The inhibition concentration (IC50) values of RTR-1 on the BGC823 cells at 24 h, 48 h and 72 h were 32.47±1.23, 15.43±0.47, and 6.40±0.03 μmol/L, respectively. The inhibition concentration (IC50) values of RTR-1 on the SGC7901 cells at 24 h, 48 h and 72 h were 32.10±1.09, 16.80±0.4, and 8.38±0.56 μmol/L, respectively. These results reveal that RTR-1 inhibits the proliferation of BGC823 and SGC7901 cells.

The cell cycle plays a crucial role in the regulation of cell growth, differentiation, senescence and apoptosis. Cell cycle progression is regulated through major checkpoints of the G1/S or G2/M phases. Additionally, the cdc2/cyclin A and cdc2/cyclin B complex promote the G2/M transition. Phosphorylation of CDK1(cdc2) on Tyr15 and Thr14 is known to be performed by the Wee1 and Myt1 protein kinases. [19] Moreover, cdc2 is phosphorylated at Tyr15 and Thr14 by Wee1 and Myt1, which induces cell cycle arrest in the G2/M phase. [20] Indeed, a significant population of the BGC823 and SGC7901 cells was arrested in the G2/M phase after treatment with RTR-1 (Figure 4A). Western blot assays also showed that the expression levels of cyclin D1, CDK1 (cdc2) and CDK4 were dramatically reduced after 24 h of RTR-1 treatment (Figure 4B and C). However, the expression levels of cyclin E and cyclin A were unchanged, and the expression levels of Myt-1, p-Weel, cyclin B1 and p-cdc2(Tyr15) increased after 24 h of RTR-1 treatment (Figure 4B and C). This observation confirms that the induction of cell cycle arrest in the G2/M phase is partially responsible for the inhibited proliferation of gastric cancer cells.

ROS could be responsible for DNA damage and, accordingly, for the activation of the G2/M DNA damage checkpoint signalling pathway. [21, 22] Indeed, we found that the level of reactive oxygen species in the BGC823 and SGC7901 cells increased significantly with RTR-1 treatment in a dose- and time-dependent manner (Figure 6A). The cell cycle checkpoints and DNA repair mechanisms constitute the first line of defence against DNA damage. Western blot analysis showed that the expression levels of p-ATM, p-chk2, p53, and p21 were increased (Figure 4B and C). In addition, we found that 5 mmol/L NAC can reverse the level of ROS production and upregulate the expression level of p-chk2 (Figure 7C). Therefore, ROS play significant roles in arresting the cell cycle in BGC823 and SGC7901 cells.

STAT3 has been shown to be involved in the processes of cell proliferation, cell differentiation and cell survival and, therefore, plays an important role in tumorigenesis, which is mediated through the regulation of various downstream target genes, including c-Myc, JunB, Mcl-1, survivin, Bcl-2, Cyclin D1, MMP-2 and vascular endothelial growth factor (VEGF). [23, 24] Previous studies have established that STAT3 may have a therapeutic effect by preventing gastric cancer, and the inhibition of activated STAT3 could reverse the resistance to chemotherapy agents of human gastric cancer cells. [25] In this study, we found that the STAT3 signalling pathway was inhibited by RTR-1 and, moreover, that the expression levels of p-STAT3 and the downstream related proteins related to it, including c-Myc, Mcl-1, Bcl-2 and Cyclin D1, were decreased. In addition, the expression level of p-STAT3 remained unchanged after NAC treatment of the BGC823 and SGC7901 cells. Thus, these data demonstrate that RTR-1 inhibits cell proliferation by promoting apoptosis through the inhibition of the STAT3 pathway in BGC823 and SGC7901 cells.

ER stress activates a set of signalling pathways collectively termed the unfolded protein response (UPR). [26] CHOP reportedly regulates energy metabolism, cellular proliferation, and differentiation. Furthermore, ER oxidase 1α (Erol-Lα), a CHOP transcriptional target, plays an essential role in inducing the oxidation of the ER lumen. [27] In addition, the CHOP-Erol-Lα pathway can induce pro-apoptotic oxidative stress in the cytoplasm. [28]The activation of PERK, generated by the dissociation of GRP78 from PERK, leads to the induction of CHOP, which plays an important role in the switch from pro-survival to pro-death signalling. [28] Additionally, it was reported that IRE1α, a rheostat capable of regulating cell fate in UPR signalling pathways, [29] can be regulated by BiP, which is combined with or separated from IRE1α. BiP inhibits IRE1α activity by binding to it in the absence of stress. [30] Our current study shows that Erol-Lα, CHOP, PERK and BiP are increased under the ER stress induced by RTR-1 in gastric cancer cells, inducing the ER stress signalling pathway in BGC823 and SGC7901 cells. Therefore, it is not surprising that the inhibition of gastric cell proliferation is also associated with the induction of ER stress with RTR-1 treatment in BGC823 and SGC7901 cells.

Apoptosis can often be induced when cancer cells are treated with agents. Recent studies showed that apoptosis was induced via three pathways, including the death receptor-mediated pathway, the mitochondria-mediated pathway and the endoplasmic reticulum-associated pathway. [31] Each of these pathways depends on the activation of Caspases. Caspases constitute a family of cysteine proteases that play essential roles in apoptosis, necrosis and inflammation. [32] Caspase3 plays an important role in the mitochondrial death pathway. Activated caspase9 can promote caspase3 activation. Then, PARP is inactivated by caspase cleavage and is regarded as a molecular marker of apoptosis. [33] The results indicate that RTR-1 can activate pro-caspase-9 and pro-caspase-3. Additionally, PARP is cleaved to induce apoptosis. These data imply that RTR-1 induces apoptosis via the mitochondria-mediated pathway in BGC823 and SGC7901 cells, and the Western blot results are consistent with those of the flow cytometric analysis.

The mitochondria-mediated apoptotic pathway is associated with the expression and activation of the Bcl-2 protein family. The Bcl-2 family contains both anti-apoptosis proteins (Bcl-2, Bcl-XL, and Mcl-1) and pro-apoptosis proteins (Bax and Bad). This family induces the release of cytochrome c in the cytoplasm. Bax is the most important member of this group and is structurally formed as either a homodimer or a heterodimer with Bcl-2. [34] Apoptosis depends on the ratio between these two proteins: it is induced by the Bax protein and inhibited by the heterodimer formed by the Bax and Bcl-2 proteins. [35] In this study, we found that with increasing concentrations of RTR-1, the expression of the anti-apoptotic Bcl-2 family of proteins Bcl-2 and Mcl-1 are decreased, while the expression of the promoting apoptosis proteins Bad and Bax are increased. Furthermore, human IAPs (at least XIAP, c-IAP1, c-IAP2, and Survivin) play essential roles in regulating apoptosis by binding to caspase pro-caspase9 to inhibit caspases [36] and block apoptosis downstream of Bax, Bik, Bak, and cytochrome c. [37-39] In addition, the IAP XIAP contains three BIR domains and a ring finger and inhibits caspase 3, caspase 7 and caspase 9. In the mitochondrial pathway, for caspase activation, XIAP, c-IAP1, and c-IAP2 bind directly to the pinnacle caspase, pro-caspase9, to prevent it from being processed and activated by cytochrome c, as is shown when caspase activation is induced by the addition of exogenous cytochrome c in both intact cells and cell extracts. [40] The results show that the expression of XIAP, c-IAP1 and c-IAP2 is increased in a dose-dependent manner. Moreover, the expression of survivin is decreased in the BGC823 cells and is maintained unchanged in the SGC7901 cells. Collectively, these data support the notion that RTR-1 inhibits cell proliferation by inducing apoptosis through the mitochondria-mediated pathway and IAP pathway in BGC823 and SGC7901 cells.

Taken together, these results demonstrate that RTR-1 inhibits cell cycle progression by the ATM–Chk2–p53–p21 pathway and induces apoptosis by activating ER stress and inhibiting the STAT3 pathway in BGC823 and SGC7901 cells (Figure 8).

## Conclusion

In summary, our findings indicate that RTR-1 inhibits the proliferation of gastric cancer (BGC823 and SGC7901) cells by blocking the cell cycle at the G2/M phase through the ATM-Chk2-p53-p21 signalling pathway and inducing cell apoptosis by inhibiting the Stat3 signalling pathway and activating the ERS signalling pathway. Thus, RTR-1 has good potential as a promising drug candidate for the treatment of gastric cancer.

## Acknowledgments

The study was supported by Science and Technology Planning Project of Guangdong Province, Grant No. 2018B030320007; Science and Technology Planning Project of Guangzhou, China, Grant/Award Number: 201607010372; and the National Natural Science Foundation of China, 81673319.

## Author Contributions

All authors contributed to data analysis, drafting or revising, the article, gave final approval of the version to be published, and agree to be accountable for all aspects of the work.

